# Methyl Halide Transferase-Based Gas Reporters for Quantification of Filamentous Bacteria in Microdroplet Emulsions

**DOI:** 10.1101/2022.12.05.519239

**Authors:** Xinhao Song, Sarah J. Kong, Seokju Seo, Ramya Ganiga Prabhakar, Yousif Shamoo

## Abstract

The application of microfluidic techniques in experimental and environmental studies is a rapidly emerging field. Water-in-oil microdroplets can serve readily as controllable micro-vessels for studies that require spatial structure. In many applications, it is useful to monitor cell growth without breaking or disrupting the microdroplets. To this end, optical reporters based on color, fluorescence, or luminescence have been developed. However, optical reporters suffer from limitations when used in microdroplets such as inaccurate readings due to strong background interference or limited sensitivity during early growth stages. In addition, optical detection is typically not amenable to filamentous or biofilm-producing organisms that have significant non-linear changes in opacity and light scattering during growth. To overcome such limitations, we show that volatile methyl halide gases produced by reporter cells expressing a methyl halide transferase (MHT) can serve as an alternative non-optical detection approach suitable for microdroplets. In this study, an MHT-labeled *Streptomyces venezuelae* reporter strain was constructed and characterized. Protocols were established for the encapsulation and incubation of *S. venezuelae* in microdroplets. We observed the complete life cycle for *S. venezuelae* including the vegetative expansion of mycelia, mycelial fragmentation, and late-stage sporulation. Methyl bromide (MeBr) production was detected by gas chromatography-mass spectrometry (GC-MS) from *S. venezuelae* gas reporters incubated in either liquid suspension or microdroplets and used to quantitatively estimate bacterial density. Overall, using MeBr production as a means of quantifying bacterial growth provided a 100-1000 fold increase in sensitivity over optical or fluorescence measurements of a comparable reporter strain expressing fluorescent proteins.

**Importance:** Quantitative measurement of bacterial growth in microdroplets *in situ* is desirable but challenging. Current optical reporter systems suffer from limitations when applied to filamentous or biofilm-producing organisms. In this study, we demonstrate that volatile methyl halide gas production can serve as a quantitative non-optical growth assay for filamentous bacteria encapsulated in microdroplets. We constructed an *S. venezuelae* gas reporter strain and observed a complete life cycle for encapsulated *S. venezuelae* in microdroplets, establishing microdroplets as an alternative growth environment for *Streptomyces spp*. that can provide spatial structure. We detected MeBr production from both liquid suspension and microdroplets with a 100-1000 fold increase in signal-to-noise ratio compared to optical assays. Importantly, we could reliably detect bacteria with densities down to 10^6^ CFU/mL. The combination of quantitative gas reporting and microdroplet systems provides a valuable approach to studying fastidious organisms that require spatial structure such as those found typically in soils.

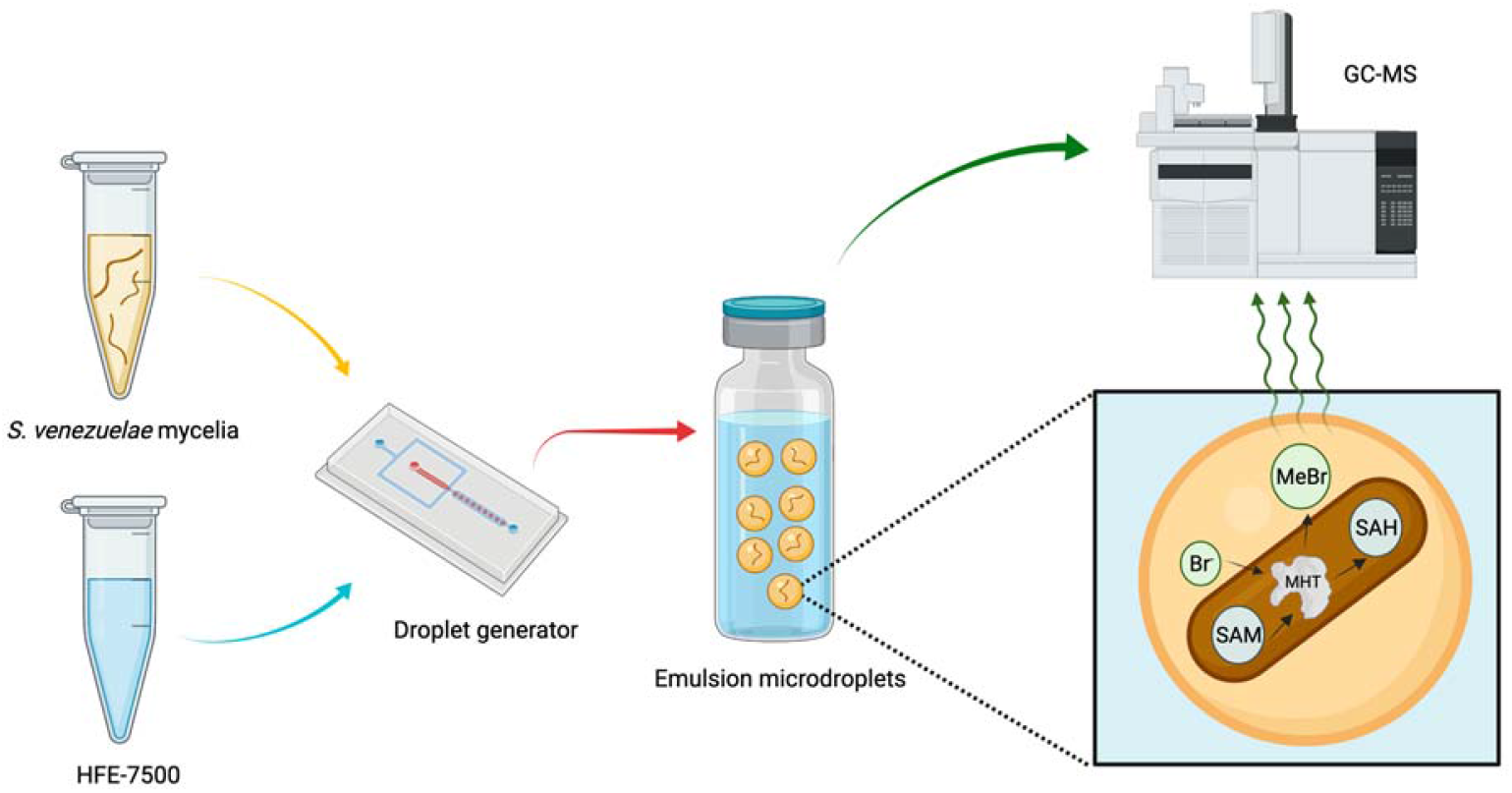

## Introduction

Application of microfluidic techniques in microbiology and environmental biology is a rapidly emerging field with substantial potential to provide new insights ^1 2^. Recent studies have used microdroplet emulsions as ultrahigh-throughput approaches to experimental evolution of enzymes ^3 4^ and microbes ^5 6 7^. Although the traditional serial transfer approach in flasks and bioreactors has been proven to be a powerful tool ^8 9^ for studies such as multicellularity ^10^, antibiotic resistance ^11 12^, and social behaviors ^13^, it suffers from several significant limitations. First, the well-mixed batch environment typically favors individuals with higher growth rates ^14^, leading to the loss of potentially relevant yet slow-growing mutants ^15^. In addition, the scalability of serial transfer studies is often limited. Compared to well-mixed batch suspensions, emulsions of homogeneous microdroplets can be used to create a compartmentalized environment in which each microdroplet can be considered an independent vessel, allowing precise control of the microenvironment and spatial structure ^16^. Importantly, such spatial segregation not only confines cells to a well-defined microenvironment and reduces competition for resources, but can also limit the diffusion of secreted molecules and privatize such “public goods” to the producers and their direct descendants within individual microdroplets ^16^, thereby favoring mutants with more offspring and allowing the rise of rare social behaviors with indirect fitness benefits ^17 18^.

Microdroplets can be manufactured over a wide range of sizes to meet the specific needs of an experimental design. For studies of prokaryotes, microdroplets ranging from ∼35 to 250 μm in diameter can be generated and can hold anywhere from one to several thousand cells ^19^. Two key factors that can affect the outcomes of experimental evolution studies in emulsions are the carrying capacity, which describes the maximum number of cells that can be supported within a single microdroplet, and the “λ value”, which describes the average number of cells in each microdroplet. Using flow-focusing microfluidic devices, the number of cells in each microdroplet typically follows Poisson distribution ^20^. Monitoring the growth of a small number of cells especially at early growth stages in real-time without disrupting the microdroplet integrity is difficult, but can provide important insights into population dynamics and cellular interactions ^21^. To this end, optical reporter systems including absorbance ^3^, fluorescence ^22 23^, and luminescence ^24^ have been developed. However, one major limitation of optical reporter systems when applied to emulsions is that the interfaces between oil and aqueous phases can interfere with the passage of light and, as a result, measurements are typically performed at single microdroplet level ^22 3^. Characterizations at a single microdroplet level are prone to stochastic perturbations and often require complex and customized experimental equipment and lengthy processing time ^25^. Moreover, optical signals can be interfered with by background noise from various sources including culture media ^26^ and pigment molecules produced naturally as secondary metabolites by some bacteria and fungi ^27^. In addition, assays using optical methods can be difficult to implement in experiments involving filamentous organisms ^28^, biofilm-producing organisms ^29^, and opaque culture media ^30^. To circumvent the aforementioned difficulties, we propose to use the production of a volatile gas ^31^ as a real-time, non-invasive, population-level growth reporting approach suitable for microbes incubated in emulsion microdroplets. This gas reporter system utilizes a *Batis maritima*-originated methyl halide transferase (MHT) that catalyzes the production of volatile methyl halide gases (MeX) from halide ions and endogenous S-adenosyl methionine (SAM). The production of MeX can then be quantified by gas chromatography-mass spectrometry (GC-MS) ^32^. Sodium bromide (NaBr) was selected as the MHT substrate as bromide ions (Br^-^) are not involved in central metabolism ^33^ and are rare in most culture media, thereby improving signal orthogonality and minimizing background noise.

As a proof-of-concept, we constructed a *Streptomyces venezuelae* gas reporter strain in which the MHT-expressing cassette was integrated into the bacterial genome. Genus *Streptomyces* belongs to phylum *Actinomycetes* and is the high GC branch of Gram-positive bacteria ^34^. *Streptomyces spp*. are crucial members of soil ecosystems and undergo a complex life cycle similar to filamentous fungi: vegetative growth by expansion of non-septic mycelia and reproduction by asexual sporulation ^35^. *Streptomyces spp*. have a remarkable potential for producing a wide array of secondary metabolites with economic or clinical significance ^36^. It is estimated that *Streptomyces spp*. have contributed around 39% of all commercial microbial metabolites ^37^ and more than two-thirds of all antibiotics ^38^. Moreover, a large number of biosynthetic gene clusters of unknown functions, termed cryptic pathways, are commonly found in genomes of *Streptomyces spp*. and are considered to be a rich reservoir for molecules with novel structures and activities ^39^. Therefore, *Streptomyces spp*. are excellent candidates for experimental evolution studies aiming at the activation of such cryptic pathways and the discovery of novel natural products for translational applications. In addition, *Streptomyces spp*. are known to form complex interspecies interaction networks in their natural habitat ^40^ presumably mediated by the secretion of diffusible secondary metabolites ^41^. The artificial spatial segregation created by microdroplets provides a unique condition for studying the evolution of social behaviors among *Streptomyces spp*. in a more controllable fashion. One technical challenge for conducting studies of *Streptomyces spp*. is that in liquid suspension, *Streptomyces spp*. typically grow as aggregated particles known as pellets ^42^, which hinders the precise quantification of bacterial density through optical methods. To solve this issue, we demonstrated that a volatile gas reporting system can serve as a quantitative non-optical method for filamentous organisms such as *Streptomyces spp*. in both liquid suspension and emulsion microdroplets.

In this study, we constructed a genome labeled *S. venezuelae* gas reporter strain (referred to as *S. venezuelae* gMHT strain hereafter) and demonstrated that MHT had activity *in vivo*. Furthermore, we explored and optimized protocols for the encapsulation and incubation of *S. venezuelae* in emulsion microdroplets. We characterized the *S. venezuelae* gMHT strain in both liquid suspension and emulsion microdroplets and showed that MeBr production was successfully detected by GC-MS in both conditions. The signal-to-noise (S/N) ratio of the gas reporting system was found to be 100-1000 fold higher than a comparable optical reporting system utilizing the constitutive expression of red fluorescent proteins in *S. venezuelae* (referred to as *S. venezuelae* gRFP strain hereafter). Moreover, good correlations between bacterial density and MeBr production were observed in both batch suspension and emulsion microdroplets, supporting the use of gas production as a real-time quantification method for filamentous organisms such as Actinomycetes.

## Results and Discussion

### Construction and characterization of a genome-integrated *S. venezuelae* methyl halide gas reporter

To demonstrate the feasibility of utilizing methyl halide gas production as a non-optical reporter system, we integrated a gas reporter system into *S. venezuelae* (ATCC 10712). *Streptomyces* was chosen in this study due to the filamentous morphology of its mycelia, which hinders precise quantifications by optical methods. The *Batis maritima* methyl halide transferase (MHT) was integrated into the genome to avoid issues such as a requirement for antibiotics and plasmid copy number instability. A ϕC31 integration system ^43^ was used to integrate the MHT cassette into the *attB* site of the *S. venezuelae* genome. It should be noted that this ϕC31 integration system integrates a spectinomycin resistance marker together with the reporter system of interest for exconjugant screening. The RFP open reading frame (ORF) on the original construct (pSC_JBEI16289) was substituted with the *mht* ORF (Fig. 1A). The vector was then transformed into donor strain *E. coli* WM6029 and introduced into wildtype *S. venezuelae* by interspecies conjugation. An RFP-labeled *S. venezuelae* strain (*S. venezuelae* gRFP strain) was also constructed using the original construct as a control. Whole-genome sequencing of *S. venezuelae* gRFP was performed and showed that the *attB* site is located at position 3,879,138 (Genbank ID: CP029197.1), which is inside of the ORF of a hypothetical pirin family protein (CDS: QES00018.1). The site of integration is consistent with a previous study in *S. ambofaciens* ATCC 23877, where the *attB* site is also located within a pirin-like gene AM23877_RS18305 (*pirA*) ^44^. Initial functional tests of the integrated MHT cassette were conducted by GC-MS. Presumptive MHT-integrated *S. venezuelae* exconjugants were incubated in CSM medium without antibiotics overnight. Mycelia were collected from the overnight culture, mixed with sodium bromide (NaBr) substrate, and subjected to GC-MS analysis. The peak of methyl bromide (MeBr) was identified at around 1 min 11 s post column injection (Fig 1B) and the position of the MeBr peak was confirmed using commercially synthesized MeBr as a standard. Antibiotic-independent constitutive expression of the MHT enzymes was observed following the genome integration of *mht*. In addition, growth curves of wildtype and labeled *S. venezuelae* were measured. Under growth conditions with no spectinomycin (Fig 1C solid lines), no significant differences were observed among the growth curves of wildtype, gRFP, and gMHT *S. venezuelae* strains, suggesting that the integration of the reporter systems did not lead to a significant metabolic burden. As expected, when spectinomycin (Spec) was supplemented (Fig 1C dashed lines), the growth of wildtype *S. venezuelae* was strongly inhibited whereas the gRFP and gMHT strains were only slightly inhibited. Using a logistic growth model, the doubling time of all strains was estimated to be ∼50 min and no significant differences were observed between wildtype and labeled *S. venezuelae* strains without spectinomycin. (Fig 1D). Taken together, we were able to integrate an MHT expression cassette into *S. venezuelae* and confirm the functionality of MHT *in vivo*. Further, although whole genome sequencing revealed that the integration event disrupted a pirin-like gene of unknown function, no significant metabolic burden in terms of growth was observed under standard growth conditions.

**Figure 1.**
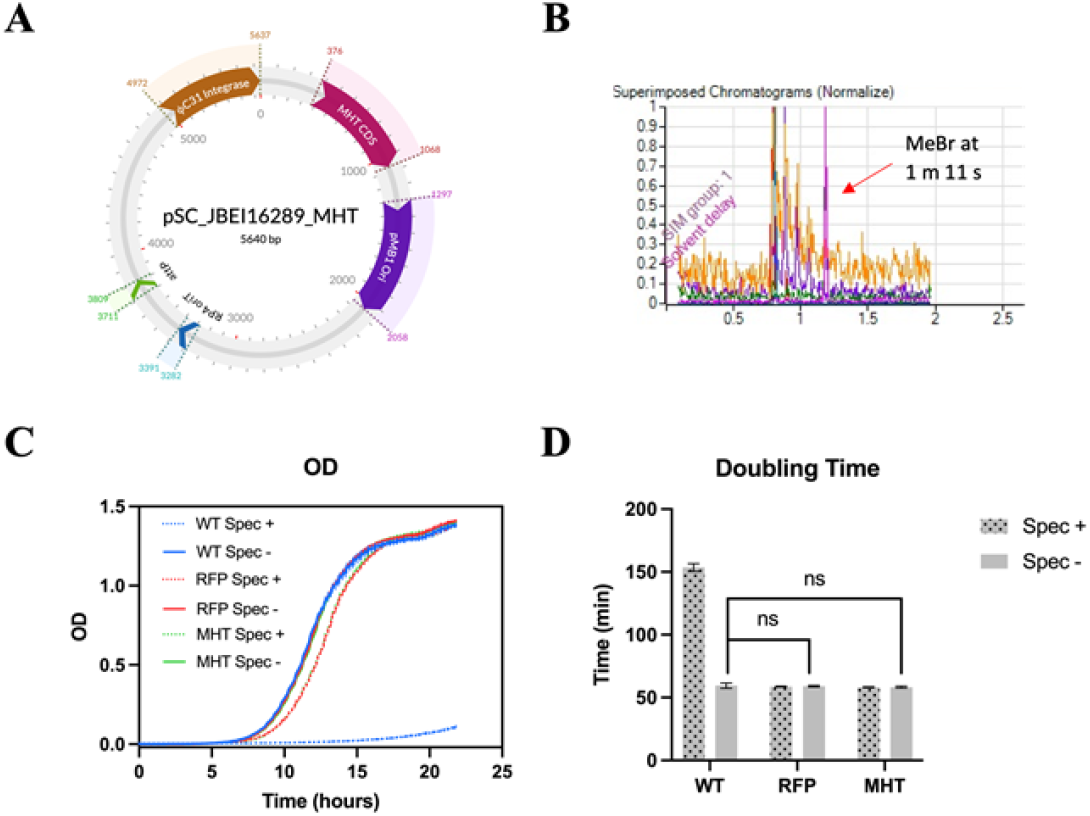
Construction and validation of *S. venezuelae* gMHT. (**A**) Plasmid map of the ϕC31-based integration vector for *Streptomyces spp*.. The RFP ORF of the original construct was replaced by the MHT ORF (red). (**B**) MeBr production of *S. venezuelae* gMHT was detected by GC-MS, indicated by the pink peak at 1 m 11 s. (**C**) Compared to wildtype *S. venezuelae* (WT), integration of reporter cassettes by the ϕC31 system did not lead to a significant metabolic burden for *S. venezuelae* gRFP and gMHT strains when spectinomycin was absent (solid lines). When spectinomycin was present (dashed lines), growth of wildtype *S. venezuelae* was strongly inhibited whereas growth of the two reporter strains was only slightly affected. (**D**) Fitting the growth curves by a logistic model showed that the integration of reporter cassettes did not significantly alter doubling when spectinomycin was absent (solid bars) compared to the wildtype *S. venezuelae*. Spec+/Spec-denote spectinomycin was present/absent, respectively.

### Characterization of methyl halide production of *S. venezuelae* gMHT strain in liquid suspension

The methyl halide production kinetics of the *S. venezuelae* gMHT strain was characterized in liquid suspension. Mycelia of *S. venezuelae* gMHT strain were collected from overnight culture and serially diluted to a final OD of 0.5 (high), 0.05 (medium), and 0.005 (low) corresponding to a CFU count of approximately 10^8^, 10^7^, and 10^6^ mL^-1^, respectively (calculated by a conversion factor of 2e8 CFU/(mL*OD)). We chose to test relatively low bacterial densities in our assays because those low densities correspond to the densities typically used in microdroplet experiments. The 24 h longitudinal readings demonstrated that the MeBr production was largely linear (with r^2^ values of 0.9737, 0.9600, and 0.9748 for OD 0.5, 0.05, and 0.005, respectively) for all three densities tested (Fig 2A). Importantly, the MeBr production of the lowest density group (OD 0.005, CFU 10^6^ mL^-1^) could be reliably detected after as early as 4 hours of incubation, exhibiting a high detection sensitivity for low bacterial densities. To further demonstrate the high detection sensitivity, we compared the signal-to-noise (S/N) ratio of the methyl halide gas production with two commonly used optical assays, namely OD and fluorescence. The *S. venezuelae* gRFP mycelium suspension was sealed in 2 mL cryogenic tubes as an approximation of GC-MS vials or dispensed into a 96-well microplate for better aeration. Except for the lowest bacterial density group (OD 0.005, CFU 10^6^ mL^-1^) at the 2 h time point, the S/N ratios of *S. venezuelae* gMHT reporter strain at all time points were 100-1000 fold higher than the S/N ratios of either OD or fluorescence measurements of *S. venezuelae* gRFP strain (Fig 2B). No significant differences in terms of S/N ratios were observed between OD and RFP fluorescence readings, although S/N ratios were improved when bacteria were incubated in 96-well microplates at 24 and 48 h time points. Importantly, both OD and fluorescence readings of medium (OD 0.05, CFU 10^7^ mL^-1^) and low (OD 0.005, CFU 10^6^ mL^-1^) density groups were indistinguishable from the background noise of blank medium and failed to detect the presence of *S. venezuelae* gRFP in liquid suspension. The inability of the optical methods to detect bacteria with low densities is presumably the result of high levels of intrinsic noise from the rich culture medium. In contrast, the background noise of MeBr gas production was much lower at all conditions tested. No endogenous enzymes have been reported to catalyze the production of methyl halide gases in *Streptomyces spp*., leading to high detection orthogonality in terms of both biological and environmental factors and an increased capability of detecting the presence of a relatively small number of bacteria at early growth stages. We further explored the feasibility of utilizing MeBr gas production as a quantification assay for bacterial density in liquid suspension. Mycelia of *S. venezuelae* gMHT strain were collected from overnight culture, normalized to an OD of 1 (CFU 2×10^8^ mL^-1^), and serially diluted from 2 to 32 folds (OD 0.03125, CFU 6.25×10^6^ mL^-1^) to produce 6 dilution groups. Normalized mycelium suspensions were loaded in GC-MS vials and MeBr production was measured after 4 and 8 hours of incubation. We demonstrated that the MeBr production readings measured at 4 and 8 h time points both followed good linear correlations with the initial bacterial densities (r^2^=0.9975 and 0.9917, respectively) and could be used as a proxy for the quantification of bacterial density (Fig 2C).

**Figure 2.**
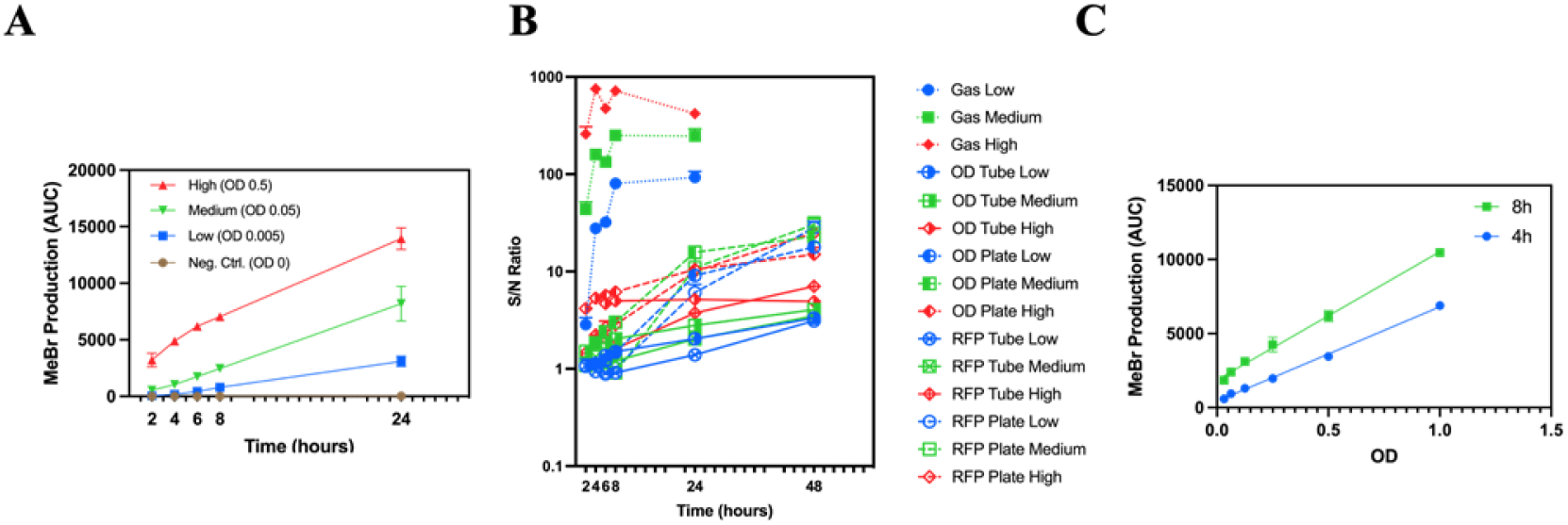
Characterization of MeBr production in liquid suspension. (**A**) Time course measurements indicated that MeBr production of *S. venezuelae* gMHT was linear over time within 24 hours in liquid suspension for all three initial bacterial densities tested: high (OD 0.5, r^2^=0.9737), medium (OD 0.05, r^2^=0.9600) and low (OD 0.005, r^2^=0.9748). (**B**) The signal-to-noise (S/N) ratios of MeBr production (dotted lines) from *S. venezuelae* gMHT were 100-1000 fold higher than measurements derived from optical signals (OD and RFP fluorescence) from *S. venezuelae* gRFP incubated in either cryogenic tubes (dashed lines) or 96-well microplates (solid lines) for all three initial bacterial densities tested: high (OD 0.5), medium (OD 0.05) and low (OD 0.005). (**C**) Measurements of MeBr production for 6 serially diluted groups ranging from OD 0.03125 (CFU 6.25×10^6^ mL^-1^) to OD 1 (CFU 2×10^8^ mL^-1^) showed that MeBr production followed good linear correlations with initial bacterial density at the 4 h (blue line, r^2^=0.9975) and 8 h (green line, r^2^=0.9917) time points.

### Characterization of growth of *S. venezuelae* encapsulated in microdroplets

To facilitate the efficient detection of methyl halide gas production, we explored the optimal encapsulation and growth parameters for the growth of *S. venezuelae* inside emulsion microdroplets. *S. venezuelae* is a filamentous bacterium and its mature vegetative mycelium fragments are typically 0.9 to 1.8 μm in diameter and up to 150 μm in length ^45^. To reduce the blocking of channels in microfluidic devices by mycelia, we used a droplet generator with 60 μm wide main channels and a 30 μm wide orifice at the flow-focusing point ^6^. Firstly, multiple combinations of aqueous and oil phase flow rates were tested and the generation of emulsion microdroplets was observed (Fig 3A). The flow rate of the aqueous phase determines the rate of water-in-oil emulsion microdroplet generation, and to maximize the generation rate, we fixed the aqueous flow rate at 50 μL/min. It was observed that the microdroplet generation was not stable when the aqueous flow rate was raised above 50 μL/min potentially due to the stratified flow effect ^46^. Therefore, to stably generate microdroplets of various sizes, we fixed the aqueous phase flow rate at 50 μL/min and adjusted the oil phase flow rate from 40 μL/min to 100 μL/min. Stable generation of highly monodispersed microdroplets was observed for all flow rate combinations tested and as expected, the diameter of microdroplets decreased as the flow rate of the oil phase increased (Fig 3A). Further, the distribution of the microdroplet diameters was estimated from 20 individual microdroplets using ImageJ. The maximum diameter achieved was 137.4±4.0 μm (with an oil phase flow rate of 40 μL/min) and the minimum diameter was 86.4±1.9 μm (with an oil phase flow rate of 100 μL/min) (Fig 3B). Soil *Streptomyces spp*. are aerobic mycelium-forming organisms ^47^, therefore microdroplets with an average diameter of approximately 110 μm (approximately 700 pL in volume) were chosen to provide ample aeration and volume for encapsulated *S. venezuelae*. Nonetheless, growth conditions in emulsion microdroplets are still drastically different from the natural soil habitat of *Streptomyces spp*. or a well-mixed liquid suspension. Therefore, experiments were conducted to evaluate how encapsulation within the microdroplet environment would affect the life cycle of *S. venezuelae* (Fig 3C). Observations were first made immediately after encapsulation, confirming the quality of microdroplets generated and a uniform presence of *S. venezuelae* mycelium fragments among microdroplets with an initial λ value of 10. At 12 h post encapsulation, initial vegetative mycelium growth could already be identified in some microdroplets. At 24 h post encapsulation, images indicated an intensive vegetative mycelium growth. Starting from 36 h until 72 h post encapsulation, sporulation at the tips of mycelium fragments (Fig 3C enlarged image on the right) and subsequent mycelium fragmentation were observed. We have shown that *Streptomyces* growing in microdroplets go through a complex developmental life cycle including the formation of mycelial mats, sporulation, and fragmentation into explorer cells ^48^. As expected, encapsulated *S. venezuelae* failed to differentiate the typical aerial hyphae structure likely due to the lack of a solid surface to provide support. Nevertheless, vegetative mycelium fragmentation and moderate sporulation were still observed, suggesting a completed life cycle for the encapsulated *S. venezuelae*. Importantly, the mycelium fragmentation we observed is consistent with a previous report of *S. lividans* pellet disintegration in liquid suspension ^49^, which makes it feasible to conduct serial evolution experiments for *Streptomyces spp*. in emulsion microdroplets. It should also be noted that heterogeneity in growth among microdroplets was observed, potentially due to variations in the number of mycelium fragments initially encapsulated or different oxygenation levels within the emulsion. Interestingly, although the size of microdroplets was relatively uniform immediately after generation, microdroplets with extensive mycelium growth shrank significantly when compared to microdroplets with less growth. One plausible explanation is that since the positions of microdroplets were relatively fixed during incubation without shaking, microdroplets closer to the surface were consistently better aerated compared to microdroplets at the bottom. Such better aeration might lead to more extensive mycelium growth of encapsulated bacteria but simultaneously more media evaporation resulting in a more significant reduction in sizes.

**Figure 3.**
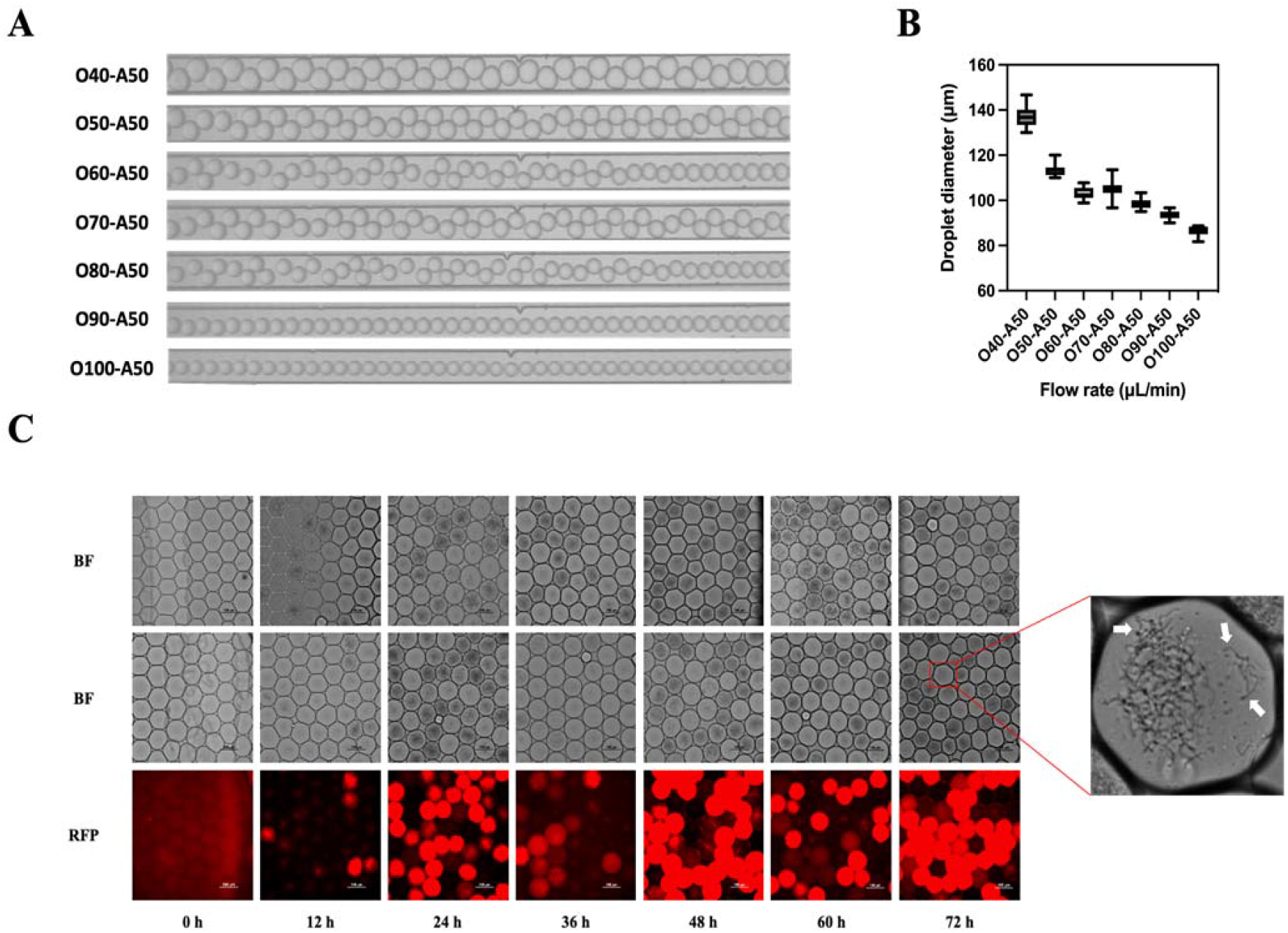
Encapsulation and growth characterization of *Streptomyces* in emulsion microdroplets. (**A**) Monodispersed emulsion microdroplets can be generated using a single device by adjusting the ratio between oil (O) and aqueous (A) phase flow rates (μL/min). The aqueous flow rate was fixed at 50 μL/min (A50) whereas the flow rates ranging from 40 μL/min (O40) to 100 μL/min (O100) were tested for the oil phase. (**B**) Diameters of microdroplets generated with various flow rate combinations were estimated from 20 individual microdroplets using ImageJ. Microdroplet size decreased as the oil phase flow rate increased and mono-dispersed microdroplets with diameters ranging from 86.4±1.9 μm (O100-A50) to 137.4±4.0 μm (O40-A50) were stably generated. (**C**) A complete life cycle of *S. venezuelae*, including vegetation growth, sporulation, and pellet fragmentation, was observed in emulsion microdroplets. The generation of mono-dispersed microdroplets and a uniform presence of mycelium fragments were confirmed immediately after encapsulation (0 h). Initial mycelium growth was observed in some microdroplets at 12 h post encapsulation and at 24 h post encapsulation, extensive mycelium growth could be observed in most microdroplets. Starting from 36 until 72 hours post encapsulation, sporulation and mycelium fragmentation were observed. The white arrows in the enlarged image on the right indicate fragmented mycelia and sporulation at the tip of such mycelia. Three representative fields are shown for each time point. Bright field channel images of *S. venezuelae* gMHT and gRFP are shown in the first and second row (BF), respectively, whereas RFP channel images of *S. venezuelae* gRFP are shown in the third row (RFP). Scale bars are at the bottom right corner of each image and denote 100 μm.

### Characterization of methyl halide production of *S. venezuelae* gMHT strain in emulsion microdroplets

Following the successful encapsulation of *S. venezuelae*, MeBr production from emulsion microdroplets was assessed. Firstly, we explored and optimized experimental factors that could potentially affect MeBr production and detection, including substrate (NaBr) concentration, sample loading volume, and microdroplet size. Substrate concentration is an important factor since an excess amount of substrates is desired to maximize gas production, but not to the point of causing growth inhibition. NaBr concentrations ranging from 50 mM to 200 mM were tested (Fig 4A). MeBr production was reduced when the NaBr concentration was either increased or decreased from 100 mM, particularly with a significant reduction (p=0.02, two-tailed unpair t-test) at 200 mM. This range is consistent with a previous study in *E. coli* in which 20 to 100 mM of NaBr was reported to be optimal for gas detection ^31^, and 100 mM was selected for all further experiments in microdroplets. The tradeoff between sample loading volume and headspace (remaining volume in GC-MS vials) also plays a crucial role in gas detection. Larger headspaces in the GC-MS vials lead to more gas accumulation and potentially higher detection sensitivity but require a smaller volume of sample. Sample loading volumes ranging from 100 μL to 900 μL were tested (Fig 4B). Interestingly, a significant improvement (p<0.05 between the 100 μL group and all other groups, two-tailed unpair t-test) in gas detection was observed for the 100 μL group. No significant differences were observed when the loading volume was more than 300 μL. Taking advantage of the fact that droplet diameter can be altered by simply adjusting the flow rate of the oil phase, we tested whether droplet size affects gas detection (Fig 4C). It was observed that smaller microdroplets (ϕ≈86 μm, oil rate=100 μL/min) led to slightly better gas detection comparing to medium (ϕ≈114 μm, oil rate=50 μL/min) and large (ϕ≈137 μm, oil rate=40 μL/min) microdroplets (p=0.0030 and 0.0073, respectively, two-tailed unpair t-test). No significant differences were observed between medium (ϕ≈114 μm) and larger (ϕ≈137 μm) microdroplets. To increase the throughput of microdroplet generation, we used a multi-channel syringe pump to generate microdroplets for multiple experimental groups in parallel and it is most convenient when the flow rates of both phases are the same. Taking this engineering factor into consideration, we selected 50 μL/min as the flow rate for both oil and aqueous phases for all further experiments.

**Figure 4.**
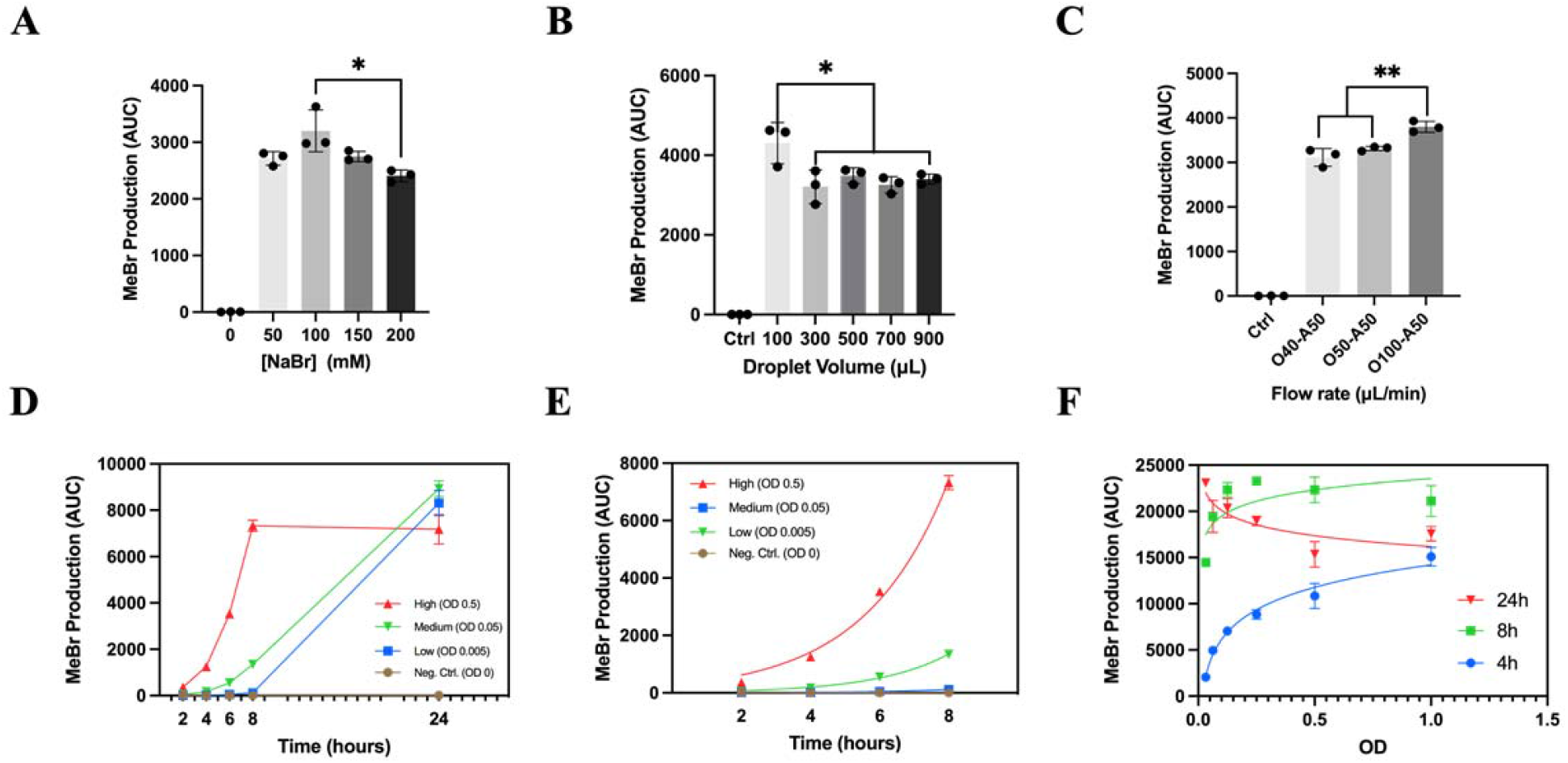
Characterization of MeBr production in emulsion microdroplets. (**A**) MeBr production was measured with substrate (NaBr) concentrations ranging from 50 to 200 mM for *S. venezuelae* gMHT encapsulated in microdroplets. A significant (p<0.05) reduction in MeBr production was observed when NaBr concentration was raised to 200 mM, with no significant differences for NaBr concentrations between 50 and 150 mM. (**B**) MeBr production was measured when microdroplets ranging from 100 μL to 900 μL in volume were loaded into GC-MS vials. A significant (p<0.05) improvement in MeBr production was observed when 100 μL samples were loaded. No significant differences were observed when the loading volume was more than 300 μL. (**C**) MeBr production was measured in microdroplets of three different sizes: large (O40-A50, ϕ≈137 μm), medium (O50-A50, ϕ≈110 μm), and small (O100-A50, ϕ≈86 μm). MeBr production was slightly improved (p<0.05) in microdroplets with the smallest size (ϕ≈86 μm). No significant difference in MeBr production was observed between microdroplets with medium (ϕ≈110 μm) and large (ϕ≈137 μm) diameters. (**D**) Time course measurements indicated that MeBr production of *S. venezuelae* gMHT encapsulated in microdroplets was non-linear over 24 hours. MeBr production of the high initial bacterial density group (OD 0.5) increased rapidly before 8 h and plateaued. MeBr production of lower initial bacterial density groups (OD 0.05 and 0.005) kept increasing within 24 hours. Notably, similar MeBr production was observed for all three groups at the 24 h time point. (**E**) MeBr production (AUC) followed an exponential increase pattern (AUC = y_0_e^kt^) over time (t) within 8 hours in microdroplets for all three initial bacterial densities tested: high (OD 0.5, r^2^ 0.9921), medium (OD 0.05, r^2^=0.9923) and low (OD 0.005, r^2^=0.8814). (**F**) MeBr production (AUC) and bacterial density (d) followed a semi-log correlation (AUC = ln(d) + b) at the 4 h time point (r^2^=0.9596). No clear correlations were observed at 8 h and 24 h time points presumably due to bacterial overgrowth.

After the values of crucial experimental parameters were determined, a longitudinal measurement of MeBr production in microdroplets was performed (Fig 4D). Mycelia of *S. venezuelae* gMHT strain were collected from overnight culture, diluted to OD 0.5, 0.05, and 0.005 (CFU approximately 10^8^, 10^7^, and 10^6^ mL^-1^, respectively) and encapsulated into emulsion microdroplets as previously described. For the high bacterial density group (OD 0.5), MeBr production increased very rapidly before 8 h and then plateaued, as indicated by similar readings at 8 and 24 h time points, whereas MeBr production of the other two groups with lower densities (OD 0.05 and 0.005) kept increasing within 24 h. Similar MeBr production was observed at the final 24 h time point for all three groups, likely due to the exhaustion of NaBr substrates. Focusing on the measurements before 8 h, MeBr production (AUC) was found to follow an exponential increase pattern (AUC = y_0_e^kt^) over time (t), with r^2^ values of 0.9921, 0.9923 and 0.8814 for OD 0.5, 0.05 and 0.005 group, respectively (Fig 4E). Next, we explored whether MeBr gas production can be used as a quantification assay for bacteria encapsulated in emulsions. Similar to the experiment in liquid suspension, mycelia of *S. venezuelae* gMHT strain were serially diluted by a factor of 2 from OD 1 (CFU 2×10^8^ mL^-1^) to OD 0.03125 (CFU 6.25×10^6^ mL^-1^) and encapsulated into emulsion microdroplets. Measurements of MeBr production were made at 4 and 8 and 24 h post encapsulation (Fig 4F) and it was observed that at 4 h post encapsulation, MeBr production (AUC) and bacterial density (d) followed a logarithmic increase correlation (AUC = ln(d) + b, r^2^ = 0.9596). No clear correlations were observed at 8 or 24 h post encapsulation, which is likely related to extensive bacterial growth and continuous MHT expression inside emulsion microdroplets. Collectively, the evidence showed that MeBr gas production could be used as a quantitative assay to accurately infer bacterial density inside emulsion microdroplets at early time points. The poor correlations observed for the 8 h and 24 h time points may be due to extensive bacterial growth in microdroplets that distorted the relative correlations of initial bacterial densities between dilution groups. In addition, it is likely that at he stationary phase, the expression of MHT is not completely halted, leading to a decoupling of bacterial density from MeBr production.

### Implications of *S. venezuelae* gas reporter in microdroplets

The concept of utilizing volatile gases as a reporting system for biological processes was initially put forward by Cheng et al. and applied to monitoring horizontal gene transfer ^31^ and gene expression ^50^ in hard-to-image soil samples. In this study, we applied the gas reporting concept to emulsion microdroplets, a condition that presents unique challenges to traditional optical assays similar to soil samples. Recent studies ^51,52^ have applied optical methods such as fluorescence-activated droplet sorting (FADS) ^22^ to investigate fluorescence signals at individual microdroplet resolution. Such an approach, however, typically requires customized optical systems and a lengthy sorting process. For example, Ran et al. ^53^ applied FADS to screen strong promoter variants that lead to a high GFP fluorescence in *S. lividans* and the detection frequency was around 10,000 microdroplets per hour. Such detection frequency could become a major time-limiting factor, especially when coupled with a much more rapid microdroplet generation process at frequencies easily exceeding 1 million per hour (Fig 3A). Moreover, for applications such as monitoring bacterial growth in real-time or investigating community dynamics, single-microdroplet resolution is unnecessary. Therefore, the gas reporting system proposed in this study serves as a non-invasive and less time-consuming assay when a quick population-wise survey is desired.

We chose to express the gas reporting system from a genome-integrated cassette instead of on plasmids to avoid two major complications. Firstly, to quantitatively evaluate bacterial density, a relatively stable spatial and temporal distribution of the reporter enzymes is crucial. Plasmid copy number is known to vary between individual cells and in different growth stages ^54^, leading to potential fluctuation in gas production. Copy number of a genome-integrated cassette, on the other hand, is definitively fixed among all individual cells and in any growth stage, providing a more stable expression for bacteria undergoing complex life cycles such as *Streptomyces* spp.. In addition, high-copy number plasmids might impose a metabolic burden on the host ^55^ and lead to a reduced growth rate. In contrast, we demonstrated that the genome integration of the reporting system led to no significant metabolic burden to the host (Fig 1C and 1D). To integrate the reporter cassette into the genome, we chose the phage integrase-mediated ϕC31 integration system ^56 43^ over a CRISPR/Cas system optimized for *Streptomyces spp*. ^57^ because no preexisting knowledge regarding the bacterial genome is required to perform the ϕC31 integration. Importantly, designing a CRISPR/Cas vector requires sequence information of the integration target region for each species to be labeled, whereas a ϕC31 integration vector containing the desired reporter cassette theoretically works in any organism with an *attB* site in its genome. The ability to label various species with one ϕC31 integration vector is particularly advantageous when studying multiple species in a community in parallel or wild isolates from environmental samples that lack prior genome information.

The gas reporter system developed in this study can be immediately applied to the quantitative evaluation of the growth profile of *Streptomyces spp*. isolates from environmental samples in emulsion microdroplets. Compared to batch suspension in test tubes, emulsion microdroplets better mimic the natural soil habitat ^31^ and can be suitable to incubate soil isolates that are slow-growing or fastidious in liquid suspension. Furthermore, emulsion microdroplets are suitable for studying secondary metabolites that are produced in the late stationary phase or during sporulation ^58^, as we were able to demonstrate a complete life cycle of *S. venezuelae* and maintain microdroplet integrity for at least 72 hours (Fig 3C). The gas reporter can also be applied to interspecies interactions or community dynamics studies in emulsion microdroplets. Using an MHT-labeled reporter as the focal species, one can study its response to various other species growing together in close proximity in microdroplets, potentially triggering the activation of cryptic pathways and the production of novel metabolites ^59^. In addition, by introducing reporters producing other volatile gases such as ethylene (C_2_H_4_) ^60^, one can monitor the population dynamics of multiple species of interest in real-time simultaneously. Particularly, thanks to the high detection sensitivity of the gas reporting system (Fig 2B), unique insights could be obtained by studying dynamics at early growth stages or behavior of minority species in a community with bacterial densities down to 10^5^ to 10^6^ CFU/mL.

To accurately quantify gas production, microdroplet samples need to be sealed in capped GC-MS vials and then incubate for 2 to 4 hours to allow a sufficient gas accumulation in the headspace (Fig 2A and 4D). Therefore, information inferred from such measurements is considered as “snapshots” from hours ago rather than the current state. In addition, although measurements can be performed for samples down to 100 μL (Fig 4B), microdroplets loaded into GC-MS vials cannot be recovered for downstream processes. In future studies, gas reporters encapsulated in microdroplets can be incubated in miniaturized bioreactor systems ^61^ with built-in gas sensors that perform headspace sampling and analysis *in situ*. Such an experiment apparatus could achieve a higher level of automation, a better temporal resolution of monitoring, and reduced sampling waste. Furthermore, although good correlations between bacterial density and gas production were observed in both liquid suspension (Fig 2C) and microdroplets (Fig 4F), mechanistic studies aiming at elucidating the distinct gas production dynamics in microdroplets are desired.

### Conclusion

In this study, we proposed the measurement of MeX gas production using GC-MS as a real-time and non-invasive growth quantification assay for filamentous bacteria in conditions where optical measurements are challenging to perform. As a proof-of-concept, we constructed a gas reporter strain in *S. venezuelae*, a model filamentous bacterium, in which the MHT cassette was integrated into the genome. Experiments in liquid suspension confirmed both the stable expression and the functionality of the MHT enzyme. MeBr production was found to be linearly correlated to the bacterial density at early time points, demonstrating the feasibility of utilizing the gas production as a growth quantification assay. Importantly, the S/N ratio of gas production signals was 100-1000 fold higher than canonical OD and fluorescence signals, suggesting a better performance for characterizing small bacterium populations at early growth stages. We then explored and optimized the encapsulation and incubation protocols of *S. venezuelae* in emulsion microdroplets and found that the microdroplet environment can support a complete life cycle of *S. venezuelae* and that growth was typically more robust than a well-mixed batch culture. Subsequently, we performed experiments encapsulating *S. venezuelae* gMHT in microdroplets. We successfully detected MeBr signals from intact microdroplets and found a semi-log correlation between MeBr production and bacterial density at an early time point. We further demonstrated that the gas reporting system can be used as a quantitative growth assay for filamentous bacteria encapsulated in microdroplets, adding a novel and practical tool that further expands the capabilities of the emulsion microdroplet platform for applications in environmental microbiology.

## Materials and Methods

### Bacterial strains and growth media

*S. venezuelae* ATCC 10712 was cultured in liquid complete supplement mixture (CSM) medium or on International *Streptomyces* Project medium 2 (ISP-2) agar (277010, BD). To make CSM medium base, 30 g of tryptic soy broth, 1.2 g of yeast extract, and 1 g of MgSO_4_ was dissolved in 1 L water and autoclaved. Upon use, 1% (v/v) of filter-sterilized 50% (w/v) glucose and 40% (w/v) maltose were supplemented to CSM base. Spectinomycin at a final concentration of 50 μg/mL was supplemented when needed. *E. coli* WM6029 was cultured in Luria Bertani (LB) medium supplemented with spectinomycin and 2,6-diaminopimelic acid (DAP) at a final concentration of 50 μg/mL and 0.1 mM, respectively. Interspecies conjugation between *E. coli* WM6029 and *S. venezuelae* was performed on AS1 agar. To make the AS1 agar base, 2.5 g of soluble starch, 1.25 g of yeast extract, 0.5 g of NaCl, and 9 g of agar were mixed in 440 mL of water and autoclaved. After autoclave, the AS1 base was cooled to ∼70 °C and mixed with 50 mL autoclaved 10% (w/v) Na_2_SO_4_ and 5 mL filter sterilized ARN amino acids mixture (1 g of alanine, 1 g of arginine, and 1 g of asparagine dissolved in 50 mL sterilized water).

### Construction of integration vector with a methyl halide transferase cassette

The *Streptomyces* integration vector ^43^ was a gift from Dr. J. Chappell (Rice University). A vector containing methyl halide transferase (MHT) coding sequence ^31^ was a gift from Dr. J. Silberg (Rice University). Gibson assembly was performed to replace the RFP coding sequence in the original *Streptomyces* integration vector with the MHT coding sequence. For Gibson assembly, both the integration vector backbone and the MHT coding sequence were PCR amplified with 20 base pair homologous overhangs on both ends. PCR products were purified and subjected to DpnI (R0176S, NEB) digestion. Gibson assembly was performed per the manufacturer’s instruction (E2611S, NEB). Gibson assembly products were transformed into NEB 5-alpha competent *E. coli* (C2987H, NEB) by heat shock. Transformed colonies were picked and the sequences of recombined vectors were confirmed by Sanger sequencing.

### Conjugation transfer of integration vector to *S. venezuelae*

The *Streptomyces* integration vector with an MHT cassette was introduced into *S. venezuelae* through *E. coli*-mediated interspecies conjugation. The conjugation donor strain *E. coli* WM6029 ^62^ was a gift from Dr. J. Chappell (Rice University). The WM6029 strain has genome-integrated *tra* locus and is methylation deficient (*dam*^-^/*dcm*^-^), allowing for efficient plasmid transfer to host *S. venezuelae* by conjugation. Firstly, the donor strain was made chemically competent following standard TSS protocol ^63^ and transformed with verified MHT integration vectors. Transformed donors were obtained and sequences of the MHT integration vector were verified by Sanger sequencing. Next, *E. coli* donors and wildtype *S. venezuelae* mycelia were collected from stationary phase cultures, resuspended in 2 mL fresh CSM media, and mixed at a 1:1 (v/v) ratio. 100 μL of the mixture was spread onto the center of AS1 ^64^ plates supplemented with 0.5 mM 2,6-diaminopimelic acid (2,6-DAP) and incubated at 30°C for 16 to 18 hours. After incubation, the lawn of *E. coli* donors was gently scraped off from the agar surface using a cell spreader, and 250 μL of an antibiotic cocktail (10 μL 30 mg/mL nalidixic acid and 20 μL 50 mg/mL apramycin mixed in 220 μL sterilized water) was applied to the agar surface. Finally, AS1 plates were incubated at 30 °C for additional 2 to 4 days until *S. venezuelae* exconjugant colonies became visible. Potential exconjugant colonies were first picked up and re-streaked onto ISP-2 plates with 50 μg/mL apramycin. Then, single colonies were inoculated into CSM media (supplemented with 5 mg/mL D-glucose and 4 mg/mL D-maltose) and passaged 3-5 rounds without antibiotics and 2,6-DAP. The presence of the integrated MHT expression cassette was confirmed by colony PCR.

### Genomic sequencing and analysis

Genomic DNA was prepared using a Microbial DNA Isolation Kit (Qiagen) per the manufacturer’s instructions. To digest bacterial cell walls and increase DNA yield, 300 μL of *Streptomyces* mycelium suspension was mixed with 50 μL 100 mg/mL lysozyme and 5 μL mutanolysin and incubated at 37 °C for 1 hour prior to DNA extraction procedures. Concentrations of the extracted DNA samples were determined using the Quant-iT PicoGreen dsDNA assay kit (P7589, Invitrogen) per manufacturer’s instructions and normalized to 2.5 ng/μL in sterilized 10 mM Tris-HCl (pH 8.0). Next, genomic DNA libraries were prepared with i5 and i7 dual index sequences attached using a plexWell 96 Library Preparation Kit (seqWell) per the manufacturer’s instruction. The genomic libraries were then sent to Genewiz for next-generation sequencing (NGS) using an Illumina HiSeq system. Sequencing was conducted in a pair-ended fashion with an average read length of 150 bp and a theoretical coverage of 100. After sequencing was completed, short-read results were aligned and analyzed using the breseq pipeline ^65^. All computational analysis was conducted on the NOTS supercomputing cluster provided by the Center for Research Computing of Rice University.

### *Streptomyces* growth curve measurement and quantification

*S. venezuelae* mycelia were collected from stationary phase culture and normalized to an OD of 0.05 using a spectrophotometer (Varian) with a cuvette of 1 cm path length. Then, 5 μL of normalized mycelium suspension was inoculated into 100 μL fresh CSM media and loaded into a 96-well microplate (Celltreat 229596). OD600 and RFP fluorescence (when applicable) of each well were measured every 10 minutes using a microplate reader (Tecan Spark) with 2.5 mm continuous shaking at 30 °C for 22 hours. Using an R package “growthcurver” (version 0.3.1), growth parameters (including growth rate, doubling rate, etc.) were fitted based on the logistic growth model. Each parameter was fitted and averaged from 3 individual replicates.

### Microfluidic device fabrication

Geometry of microfluidic devices was designed using CAD software and then fabricated onto silicon wafers using photolithography with a thin layer of SU-8 2075 photoresist material (Kayaku Advanced Materials) per the manufacturer’s instructions. Liquid polydimethylsiloxane (PDMS) was mixed with 10% (wt/wt) curing agent Sylgard 184 (Electron Microscopy Sciences), degassed in a vacuum chamber for 10 min, and then applied onto the surface of silicon wafer molds. After 1 to 2 hours of incubation at 80 °C, solidified PDMS was peeled off from the silicon wafer and cut into individual chips per design. Inlet and outlet channels were created using a 0.5 mm core sampling puncher (Electron Microscopy Sciences) under a microscope. Individual chips were then immersed into methanol and sonicated for 8 min to remove debris in channels. Finally, PDMS blocks were bound to microscope glass slides after 2 min plasma activation with ozone at 100 mTorr in a plasma cleaner (PDC-001, Harrick Plasma). Devices were incubated for an additional 16 to 18 hours for complete binding ^66^.

### Encapsulation of *S. venezuelae* into emulsion microdroplets

*S. venezuelae* mycelia were collected from stationary phase culture and resuspended in fresh CSM media as the aqueous phase. 100 mM sodium bromide (NaBr) was supplemented to the aqueous phase as the substrate for MHT when necessary. HFE-7500 (Novec, 3M) was supplemented with 1.5% (vol/vol) surfactant Pico-Surf 1 (Sphere Fluidics) and employed as the oil phase. Both phases were injected into the droplet generator device through designated inlets at a flow rate of 50 μL/min using a multichannel syringe pump (NE-1200-US, New Era Pump Systems), and emulsion microdroplets of an average diameter of approximately 110 μm were generated. Droplets were collected into 50 mL centrifuge tubes for incubation or further characterization.

### Quantification of methyl halide production by GC-MS

MHT labeled *S. venezuelae* mycelia were collected from stationary phase culture, diluted to desired bacterial density with fresh CSM media, and supplemented with 100 mM sodium bromide (NaBr). For quantification of gas production in liquid suspension, 1 mL of bacterium suspension was transferred into 2 mL glass vials (AR0-37K0-13, Phenomenex) and sealed with designated caps (AR0-5760-13, Phenomenex). For quantification of gas production in emulsion microdroplets, the bacterium suspension with NaBr was used as the aqueous phase for droplet generation. Then, 500 μL of the microdroplet emulsion was transferred into GC-MS vials. Vials were incubated at 30 °C without shaking for 2 to 4 hours to allow for initial gas accumulation. At each time point, MeBr production was measured with an Agilent GC-MS system (8890 GC system 8890 coupled with 5977B GC/MSD system with a 7693A autosampler). MeBr ions (m/z = 94.94) were detected at a retention time of 1.18 min as determined previously by a pure MeBr standard. Carbon dioxide (CO_2_) ions (m/z = 44.01) were also detected at a retention time of 0.81 min as a control. MeBr production was quantified by the area under curve (AUC) metric by Agilent MassHunter Quantitative Analysis software with a qualifier ratio range between 78.1 and 117.1.

### Microscopy of *S. venezuelae* in emulsion microdroplets

To visualize the encapsulated streptomycetes, approximately 1 μL of emulsion droplets were collected into a rectangular capillary glass tube with an inner diameter of 100 μm (VitroTubes, 5015-050). The capillary tube containing microdroplets was placed on a microscope slide (1 mm thick) and then both ends of the capillary tube were sealed with Lubriseal stopcock grease (8690-B20, Thomas Scientific) to avoid excessive microdroplet movement during imaging. The slide was loaded onto a Nikon Eclipse Ti2 fluorescence microscope for imaging in both the bright field channel (optimal exposure time) and RFP channel (excitation = 560 nm, emission = 635 nm, exposure time = 500 ms).

### Real-time monitoring of microdroplet generation and quantification of microdroplet diameter

Droplet generator device was loaded onto a Nikon Eclipse Ti2 fluorescence microscope and microdroplets generated were monitored in real-time using a Phantom VEO highspeed camera (Ametek) and images were taken at 5200 fps with 190 μs exposure time. Diameters of microdroplets were manually measured using ImageJ software with a conversion factor of 0.3 pixel/μm. The average and standard deviation of each flow rate combination were calculated from 20 independent measurements.

### Data availability

All relevant data, including original microscope images, GC-MS and microplate reader raw outputs can be provided upon request.

## Acknowledgments

The authors would like to acknowledge Dr. James Chappell of Rice University for the *Streptomyces* integration vector pSC_JBEI16289 and the conjugation donor strain *E. coli* WM6029. The authors would like to acknowledge Dr. Jonathan Silberg of Rice University for the MHT-containing vector. The authors would like to acknowledge Dr. Carrie Masiello of Rice University for her help on GC-MS experiments. This work was supported by the National Institutes of Health, National Institute of Allergy and Infectious Diseases grant R01A1080714 to Y.S.

